# Fast and inexpensive detection of bacterial viability and drug effectiveness through metabolic monitoring

**DOI:** 10.1101/042499

**Authors:** Sondos Ayyash, Wen-I Wu, P.Ravi Selvaganapathy

## Abstract

Conventional methods for the detection of bacterial infection such as DNA or immunoassays are either expensive, time consuming, or not definitive; thus may not provide all the information sought by the medical professionals. In particular, it is difficult to obtain information about viability or drug effectiveness, which are crucial to formulate a treatment. Bacterial culture test is the “gold standard” because it is inexpensive and does not require extensive sample preparation, and most importantly, provides all the necessary information sought by healthcare professionals, such as bacterial presence, viability and drug effectiveness. These conventional culture methods, however, have a long turnaround time: anywhere between 1 day to 4 weeks. Here, we solve this problem by monitoring the growth of bacteria in thousands of nanowells simultaneously to identify its presence in the sample and its viability, faster. The segmentation of a sample with low bacterial concentration into thousands of nanoliter wells digitizes the samples and increases the effective concentration in those wells that contain bacteria. We monitor the metabolism of aerobic bacteria by using an oxygen sensitive fluorophore, ruthenium tris (2,2’-diprydl) dichloride hexahydrate (RTDP) that allows us to monitor the dissolved oxygen concentration in the nanowells. Using *E.Coli* K12 as a model pathogen, we demonstrate that the detection time of *E.coli* can be as fast as 35-60 minutes with sample concentrations varying from 10^4^(62 minutes for detection), 10^6^ (42 minutes) and 10^8^ cells/mL (38 minutes). More importantly, we also demonstrate that reducing the well size can reduce the time of detection. Finally we show that drug effectiveness information can be obtained in this format by loading the wells with the drug and monitoring the metabolism of the bacteria. The method that we have developed is low cost, simple, requires minimal sample preparation and can potentially be used with a wide variety of samples in resource poor setting to detect bacterial infections such as Tuberculosis.

## 1. INTRODUCTION

Infectious diseases caused by protozoa, bacteria, fungi and viruses, are a leading cause of human suffering globally. According to World health Organization, infectious diseases claim 16.7 million lives worldwide each year ^1^. Annually, an estimated 16 percent of all deaths worldwide result from infectious diseases ^2^. Bacterial infectious diseases are responsible for a significant portion of these deaths. Some of the deadliest bacterial diseases include tuberculosis (1.5 million deaths worldwide in 2013) ^3^, syphilis (113, 000 deaths worldwide in 2010) ^4^ and streptococcus (826,000 deaths worldwide) ^5^ just to name a few.

Despite being curable and the availability of medicines for these diseases, the global morbidity and mortality rates remain high^6^. It should be noted that while treatment for many of these conditions is inexpensive, its diagnosis and identification of a suitable treatment is not. For instance, although tuberculosis can be treated and cured, there still remains a high rate of TB related deaths worldwide. WHO reports that over 95% of Tuberculosis deaths occur in low-and middle-income countries ^5^. Thus, the development of an inexpensive diagnostic technique is key to addressing the number of cases of reported deaths caused by bacterial infectious diseases in resource poor settings.

Bacterial detection methods can have far reaching applications beyond diagnosis, such as monitoring contamination in food and water. Thus, the identification of microbial contamination on meat^7,8^microbial pathogens in water^9^, improving patient care and preventing the spread of disease ^10^^-^^12^, are all areas that would benefit from a fast, and low-cost bacterial detection method.

The current “gold-standard” for bacterial disease diagnostics^13^ is the bacterial culture test, a 120-year-old method^14^. This technique is still used in many resource poor settings for the diagnosis of infectious diseases, as it is widely available, easy to handle, and inexpensive^14^. Using a selective media that promotes the growth of specific bacteria, the conventional culture test method is able to provide information on the presence, viability and drug effectiveness of that bacteria ^15^. All of this information is crucial for healthcare professionals in diagnosis and in formulating a treatment. However, it takes a long time, as visualization of a colony by eye is the threshold for detection. For instance, it is known to take 2-6 weeks for visual detection of the presence of tuberculosis colonies on the surface of a culture plate.^16^ Even *E.Coli*, a fast growing bacterium, requires a day of culture. Due to this disadvantage, molecular diagnostic methods including DNA biosensors, DNA chips,^17^ immunoassays (ELISA), and nucleic acid assays (PCR)^18^ have been developed. They are faster than the culture test but are either expensive^19^, require extensive sample preparation^20^or may not provide all the information such as viability^21,22,23^and drug effectiveness^19^sought by doctors in diagnosing a disease and formulating a treatment. Another kind of bacterial detection based on biological detection by optical oxygen sensing, known as fluorescence-based respirometric screening technology (RST)^24,25^ allows convenient high-throughput analysis of oxygen consumption by cells, however it is not able to perform rapid diagnostics especially when the concentration levels of bacteria are very small. For instance, it still takes 6-8 hours for detection of 10^4^ cells/mL of *E.Coli* using RST while higher concentrations such as 10^6^ cells/mL can be detected in ~2 hrs.^26^

Here, we use metabolic monitoring of the growth of bacteria in nanoliter well arrays to increase the speed of detection of bacteria, its viability and its drug effectiveness. We demonstrate rapid detection of the growth (~ 1 hr for 10^4^ cells/mL) and show that detection is faster when nanowells are smaller. We also demonstrate that minimal sample preparation is required for this method making it suitable for resource poor settings. This method could be a viable alternative to the current culture method and could be easily implemented in a wide variety of settings.

## 2. Working Principle

At its fundamental level bacterial culture is a simple yet robust method to identify that a particular organism is alive (viable) and to visualize it to the naked eye through amplification (colony growth). Our visual resolution then determines the smallest colony that we can see and hence the time for detection of growth.

However, there are other methods that one could use to determine viability and growth. Any organism that is alive will consume nutrients and excrete waste. Thus, by measuring the material that is consumed or excreted, one could have an early indicator of the viability of the bacteria even before it has divided and grown sufficiently to be visually noticed. This is the principle behind the metabolic monitoring of culture as a generic method to measure the state of health of the organism. Specificity is provided by the use of selective growth media that only allows the growth of specific bacteria. An example of selective media is Middlebrook broth mixed with antibiotics that is used to kill all other bacteria other than mycobacteria and is used in the detection of tuberculosis.^27^

In this paper, we devise a method to detect the bacteria faster by measuring oxygen consumption (metabolic marker). The device consists of an array of nanoliter wells, which is fabricated using softlithography as shown in figure 1a. The inside surface of the wells are made hydrophilic while the top surface is made hydrophobic. Due to this configuration in surface properties, a sample is dispensed and spread on the device will quickly fill the wells, automatically partitioning the sample into thousands of equi-sized nanoliter volumes.

**Figure 1.**
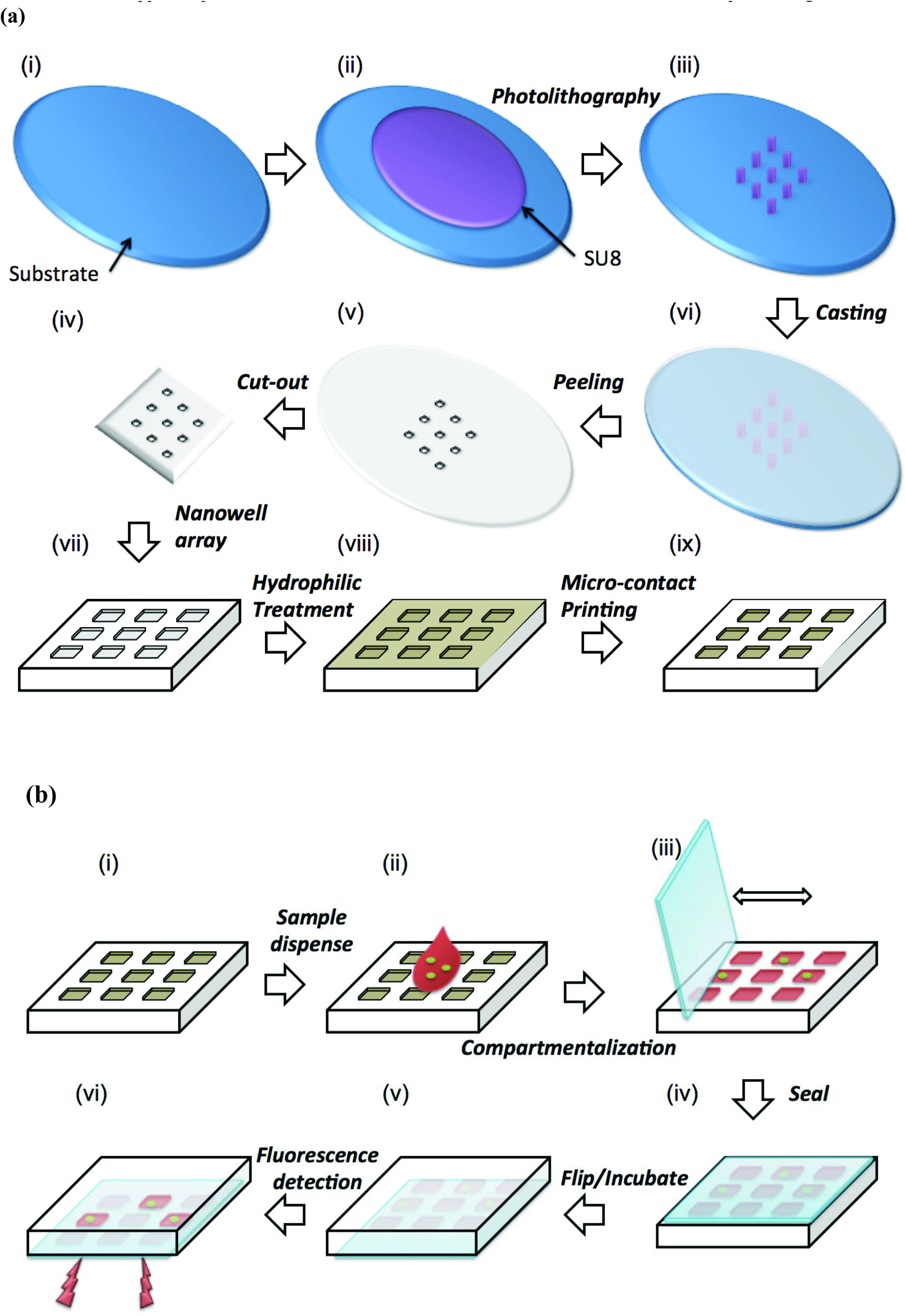
Schematic representation of the. (a) soft lithography and surface modification process to fabricate the device and (b) sequence of operation of the device

The sample is mixed with an optical, oxygen quenching fluorophore (RTDP) as well as selective media ^28^ that facilitates growth of only the specific bacteria of interest. This is accomplished prior to dispensing it onto the surface of the device containing an array of nanowells (figure 1b). A glass slide is then swiped across the surface (like a squeegee), which allows the bacterial solution to deposit and compartmentalize into the specific wells (figure 1b).

Once compartmentalization of liquids within the defined wells is completed, a glass slide seals the filled nanowells to prevent evaporation of the sample (figure 1b). The sealed device is then imaged through a fluorescent microscope or microarray scanner to measure the fluorescent intensity (metabolic measure of oxygen) changes over time (figure 1b). Since bacteria are aerobic, they consume oxygen that is present in the medium during metabolism and depletes the oxygen in the surrounding environment, producing fluorescence. The separation distance between the nanowells is determined by the permeability of the material to oxygen over the duration of the detection. Drug effectiveness can also be probed by adding the appropriate drug to the broth and measuring growth or lack of it through fluorescence.

The segmentation of the sample into thousands of individual nanoliter wells digitizes the sample and has some benefits. Consider a typical sample that is several hundred nanoliters in volume contain a certain concentration of bacteria. At high concentrations, the concentration of bacteria in each well is the same as the overall concentration of the sample. However, at low concentrations, if the sample were segmented into thousands of smaller volumes then some of the wells will contain the bacteria of interest while others will not. The process of digitization of the samples creates some wells where the local concentration of bacteria is several times higher than the overall concentration in the sample and others where there would be no bacteria. Therefore, by digitizing the sample and measuring the metabolic markers in each of the wells simultaneously, one could arrive at detection of the bacteria much faster than in a larger homogeneous sample. This is the working principle behind our fast metabolic monitoring of bacteria.

A similar segmentation can be achieved in a digital droplet microfluidic platform ^29^. However it requires sophisticated instrumentation for droplet generation compared to the method here and may not be suitable for resource poor settings. Furthermore, the high permeability of the oil barrier to gases precludes it from use when the metabolic marker is oxygen.

Large scale arrays of nanowells have been used in the past for single cell imaging and analysis^30^ and analyzing cytoplasmic contents of individual cells^31^. Recently, it has been used to perform end-point ELISA type immunoassays in the nanowell format for identification of mycobacteria^32^. However, ELISA requires extensive sample preparation for analysis, which is not only labor intensive but also a time-consuming. Also, endpoint assay typically requires that the assay duration be set for the minimum detectable concentration, as the concentration present in the sample is not known apriori.

Here we report a generic method that uses nanoliter well arrays to perform real-time measurement of oxygen as a metabolic marker to detect viability, growth and drug effectiveness of bacteria. The method requires very simple sample preparation that is suitable for resource poor settings. Real-time measurements allow faster detection of growth especially when the concentrations are higher than a single bacteria/well. The instrumentation required for imaging and fluorescent measurements can also be made simpler and cost-effective using some of the recent developments in low cost optics^33^. In combination, this approach provides an effective method for fast culture based detection in resource poor settings.

## 3. Materials and Methods

### Materials

Silicon wafers (mechanical grade, 3”, 500 μm thick) were purchased from University Wafer Co., USA. Negative photoresist Su8-100 and developer were acquired from Microchem Co., USA. PDMS polymer was purchased from Dow Corning Co., Canada. The powder of RTDP (ruthenium tris (2,2’-dipyridyl) dichloride hexahydrate) was obtained from Sigma-Aldrich (#224758). Liquid microbial growth medium, Luria Broth (LB) purchased from Sigma-Aldrich (#L2542). The hydrophilic surface modification agent (N-Wet 410) was gifted by Enroute Interfaces Inc., and one part of the n-Wet 410 solution was diluted with 9 parts of toluene giving a final concentration of 10%V/V. Glass slides (Corning, 75×25 × 1mm) were used to sandwich our device. A commercial hydrophobic surface modification agent, Aquapel was obtained from Aquapel^®^ Glass Treatment, and was used to make glass slides hydrophobic. Methylcellulose MC (M-0262) viscosity of 2% aqueous solution at 20°C, 400 centipoises was obtained from Sigma.

### Microfabrication of nanowell array

SU-8 100 photoresist is spun on a Silicon wafer at 1,700 rpm to produce a layer of 100-μm thickness (figure 1a-ii). This layer was patterned using photolithography and produced square or circular shapes of sizes that varied from 100-1000 *μ*m to produce the mold for nanowell devices (figure 1a-iii). Next, polydimethylsiloxane (PDMS) prepolymer (base: curing agent = 10:1) was cast on the mold, cured and peeled off to replicate the pattern of the mold forming an array of nanoliter wells as shown in figure 1a-iv. Next, the cast and crosslinked PDMS elastomer is peeled off and cut to the required shape. Subsequently, the device is immersed and left overnight into the n-Wet 410 solution to modify the entire surface of the PDMS to a hydrophilic state (figure 1a-viii). Finally, the top surface of the device is microcontact stamped with a thin layer of PDMS prepolymer and cross linked in order to make only that surface hydrophobic (figure 1a-ix).

### Experimental procedure

During experiments 0.01 mL sample solution containing bacteria is dispensed on the surface (figure 1b-ii). Next, a glass slide is used as a squeegee to move the liquid around on the surface. The sample liquid automatically enters and compartmentalize into the nanowells as it is moved around (figure 1b-iii). The device is then sealed with a glass slide coated using a commercial hydrophobic surface modification agent (Aquapel®) and imaged under a microscope to measure the fluorescence intensity of the oxygen quenching fluorophore. A confocal microscope was used throughout the experiments to capture images, whereby the *E.coli* is located on a specific plane, and thus, the fluorescence of the RTDP is obtained along the same location for the remainder of the experiment. Each experiment was repeated for a minimum of three times; however, the majority of experiments were repeated nearly six times. Once experiments were completed, offline image analysis of the fluorescent intensity within the wells was calculated by using ImageJ (NIH, rsb.info.nih.gov/ij).

### Data Analysis

The intensity was normalized to the initial intensity in the well for reporting purposes. For all the experiments, the initial intensity of the well was subtracted from the current intensity to obtain the normalized increase in intensity at that time point. In one case where the changes in intensity in response to the presence and absence of the drug is measured, the normalization was done by dividing the current intensity by the original intensity at t=0, in order to better represent the data.

### Device dimensions

We have developed a microfluidic device that is 1-2cm^2^ in size composed of an array of nanowells with diameters (100-1000*μ*m), shape (circular, square), volume (1nL-100nL) and a depth of 100 *μ*m. The dimensions of the *E.coli* bacterium are 0.5 × 2μm, and thus the well size was selected to be large enough to have enough nutrients for a growing population, but small enough for fast detection rates.

### Artificial sputum

The sputum was prepared by mixing 2%(w/v) methylcellulose (Sigma M-0262) in 1000mL of DI water ^34^, which resulted in a viscosity of 0.4 pa-s at 20°C^35^. This viscosity matched that of mucoid, which was reported by Maria Teresa Lopez-Vidriero et al^36^ to be 0.42 pa-s.

### Bacterial strains and growth conditions

E.coli strain (OP50 and K12) was used for this study. Cells were grown aerobically at 37 °C in LB medium (10 g tryptone, 10 g NaCl, 5 g Yeast extract per liter). Bacterial cells in stationary phase were harvested after overnight incubation.

## 4. Results & Discussion

### 4.1. Sample dispensing and compartmentalization

The sample dispensing technique was tested on a range of different nanowell sizes to determine which among them filled with ease. Various sized nanowells in the range of 1nL (35 × 35 nanowells; 1225 nanowells in total) to 100nL (7 × 7 nanowells; 49 nanowells in total) were fabricated, with a device area of 7.5 × 7.5mm. The nanowells were either square or circular in crossection and their feature size ranged between 100 to 1000*μ*m: dimensions of: 225 *μ*m (5 nL), 265 *μ*m (7 nL), 340 *μ*m (11.5nL), 360 *μ*m (13 nL), 500 *μ*m (25nL), and 1000 *μ*m (100nL). The depth of the nanowells were kept constant at 100 *μ*m. A single drop of liquid was portioned into hundreds of nanoliter-sized wells by dragging it along a surface with a hydrophobic coverslip. It is brushed onto the surface of the elastomer in the one direction (sideways) and then swiped back in the opposite direction. With each swipe, a certain percentage of wells are filled. The process of filling of the wells is demonstrated in figure 2a. Here, nanowell arrays with 100*μ*m (1nL) dimensions were filled with DI water mixed with methylene blue dye for visualization.

**Figure 2.**
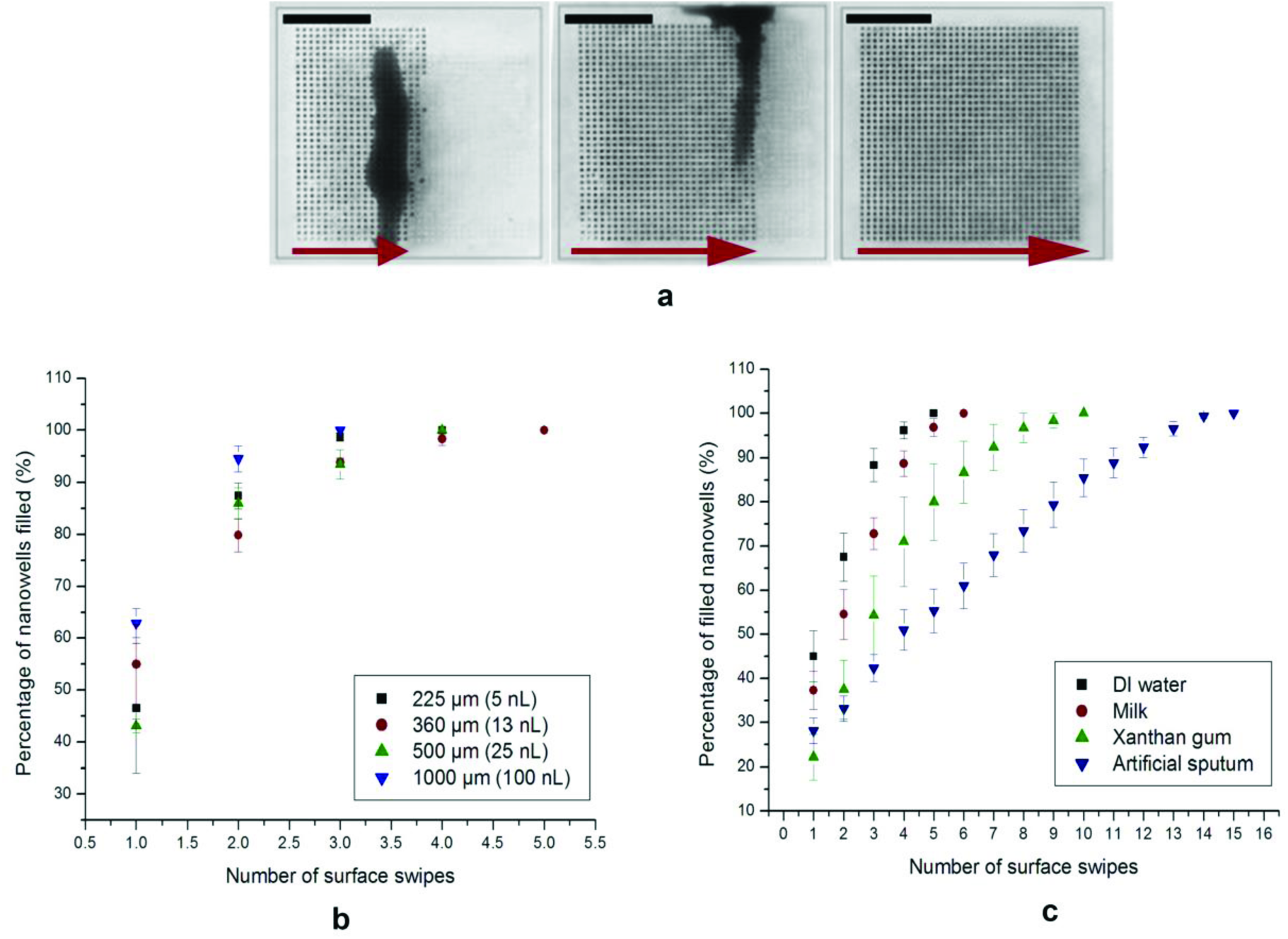
**a)** Optical images of progressive filling of an array of 1nL volumes with dimensions: l00*μ*m × l00μmxl00μm. A mixture of DI water and methylene blue is dispensed on the surface, whereby a glass coverslip is used as a squeegee to push the liquid from left to right until the entire matrix is filled. Scale bars, 2.5 mm. **b)** Percentage of nanowells of different volumes filled per swipe (with a coverslip) using a mixed solution of de-ionized water and methylene blue **c)** Percentage of wells filled with each swipe, of a 1nL array, with solutions of different viscosities: DI water (0.001 Pa-s), milk (0.003 Pa-s), xanthan gum solution (non-Newtonian fluid), and artificial sputum (0.4 Pa-s).

Various well sizes were tested and the number of swipes required to completely fill all the wells in a particular design has been plotted in figure 2b (*n=6*). The sample liquid used was DI water with a viscosity of 0.001 Pa-s. The results indicate that the wells were completely filled in all the designs with a maximum of 5 swipes. Since each of the swipes takes ~ 10 sec, the entire device could be filled in under a minute with minimal instrumentation or technique. It was evident from the results shown in Figure 2b, that the larger sized nanowell (100nL-S/V ratio of 0.024) filled slightly faster than the smaller sized nanowell array. The percentage of the wells filled were significantly different after one swipe and became nearly the same after multiple swipes. Filling the nanowells becomes a difficult task at much smaller scales, and evaporation becomes significant at 25 fL as reported by Taylor and Walt.^37^ The difference in filling between the different sized wells in the range tested was not statistically significant (p>0.5). Therefore, regardless of the size of the well, the array was filled in 3-5 swipes.

The ability of the solution to compartmentalize within the nanowell array was also characterized using liquids of different viscosities ranging from water (0.001 Pa-s at 20°C) to high viscosity shear thinning liquids such as xanthan gum (59.96% water/40% glycerol/0.04% xanthan gum)^38^, protein rich solutions such as milk (viscosity of 0.003 Pa-s at 20°C) and artificial sputum (0.4 Pa-s at 20°C)^34^. These tests were conducted to mimic real samples such as blood (protein rich) or sputum (high viscosity) and their effect on compartmentalization in our device. The experimental results as shown in figure 2c demonstrated that all the types of fluids used were able to fill the nanowells within reasonable number of swipes. However, fluids with lower viscosities (DI water) compartmentalize and fill the entire array of nanowells faster (within 5 swipes) than fluids with higher viscosities (i.e. artificial sputum - 15 swipes). The artificial sputum had nearly 400 times higher viscosity to DI water and filled only 50% of the wells at 5 swipes. This experiment demonstrates that viscosity of a solution is an important factor in the compartmentalization process. Nevertheless, since each swipe takes only ~ 10 sec, the extra time required (~100 sec) is reasonable and can easily be accommodated within the sample preparation process. Protein rich solutions such as milk with viscosities similar to water - 0.003 Pa-s, required an additional swipe compared to DI water. This result shows that high protein loading of the sample did not affect the compartmentalization and was not a significant factor in the filling of the nanowells as compared to viscosity. xanthan gum diluted in water (59.96% water/40% glycerol/0.04% xanthan gum)^38^,was used as a viscoelastic, shear thinning, blood analogue. It required 10 swipes to completely fill a nanowell array. This set of experiments demonstrated that viscosity of the sample solution is the key determinant in filling of the nanowells. It also showed that higher viscosity samples require higher number of swipes to fill. Finally, it demonstrated that a wide variety of sample types could be compartmentalized within a short time frame and without elaborate setup using our nanowell array.

### 4.2. Distribution of bacteria

The distribution of the bacterial cells within an array of nanowells after dispensing was measured for various concentration of sample solution in order to identify if it is uniform. The sample solution used was Luria Broth (LB) media spiked with GFP expressing *E.Coli* with concentrations of from 10^4^ to 10^8^ cells/mL. This sample was dispersed into a nanowell array that had well volumes of 1 nL. A uniform and complete dispersion of the sample into the wells will then produce a population of 100 cells/well at 10^8^ cells/mL and 1 cell/well at 10^6^ cells/mL. For a concentration of 10^4^ cells/mL, it is expected that about 1 out of 100 wells will contain bacterial cells. Three sets of dispensing experiments were performed on separate devices and 11 wells at random were measured in each of the devices. For the low concentration samples, 100 wells were measured. Bacterial population in each well was counted manually by taking confocal images at various depths in the nanowells. Figure 3a shows one such composite image where the GFP expressing bacteria was imaged as green dots inside the well. The oxygen sensitive fluorophore (RTDP) dispensed into the wells can be simultaneously imaged at a different wavelength (red). At the sample concentration of 10^8^cells/mL, individual wells were found to contain ~48 cells/well with a standard deviation of 10. Sample concentration of 10^6^ cells/mL resulted in 4.5 cells/well with a standard deviation of 2.2. Finally, at a concentration of 10^4^ cells/mL the bacterium was scattered and scarce, only to have 1 bacterium present in certain wells while many others contained no bacterium at all. The mean distribution of a concentration of 10^4^cells/mL was found to be 1 cell each in six separate wells out of 100 wells (6/100), with a standard deviation of 2.0. The populations at various concentrations of sample seem to correspond to their expected values to an order of magnitude. Distribution of the bacteria appeared to be uniform. Although a confocal microscope is used here to image and count the bacteria (1 um) at various levels within the nanowell for characterization purposes, the use of a metabolic marker which is distributed throughout the well, ensures that a low resolution CCD imager can be used to measure intensity of the nanowells (~ 50 μm in size) and through it identify the presence or absence of bacteria.

### 4.3. Calibration of oxygen sensing

The calibration of fluorescent sensing of oxygen concentration was carried out using samples prepared to have a known concentration of oxygen in them. First, an oxygen scavenger sodium sulphite, 5g/L was dissolved in LB media along with the fluorophore RTDP (0.3mg/mL) to produce a sample with depleted of oxygen levels. Dissolved oxygen was measured using a commercial optical oxygen sensor (YSI 550A, YSI Environmental, Yellow Springs, Ohio) which yielded a solution with an oxygen concentration of 0 mg/L. A second sample was prepared by bubbling oxygen gas into the LB media, mixed with RTDP, to produce an oxygen concentration of 15.6 mg/L. Finally, LB media exposed to ambient atmospheric condition was also tested. These samples were dispensed onto the nanowell array and intensity of fluorescence in the wells was observed over one hour as shown in figure 3b. Each condition was repeated three different times, and 14 wells were analyzed and processed for each experiment.

**Figure 3.**
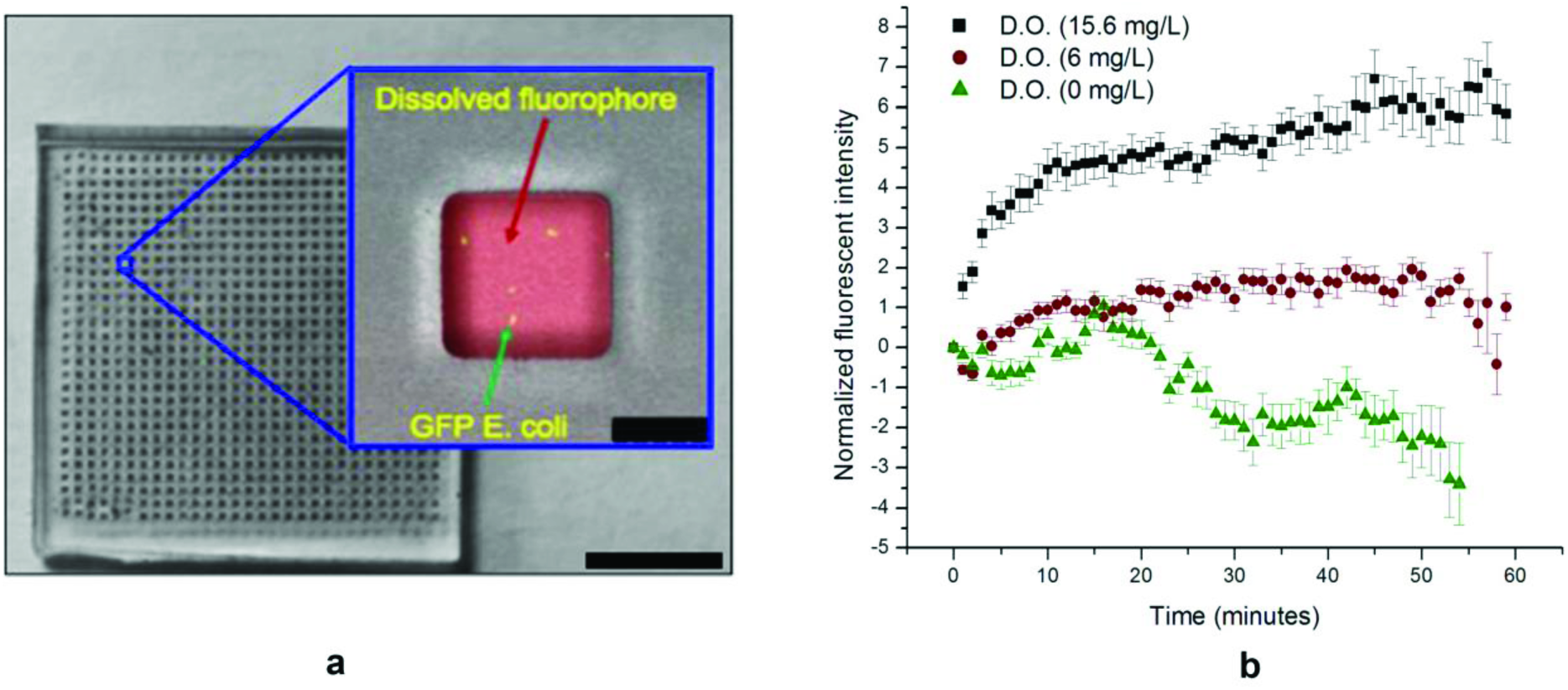
**a)**Optical image of the nanowell array. The inset shows fluorescent image of a single well with both GFP expressing bacteria (green) and the oxygen sensitive fluorophore (red). The oxygen sensitive fluorophore is dispersed throughout the entire well and indicates the oxygen concentration in the entire well. The wells volume is 1 nL. The GFP and RTDP fluorophore were imaged separately using a different combination of excitation and emission filters, and then overlaid to give a complete image with bright-field. Scale bar of device, 25.5 mm. Scale bar of zoomed in well, 50 μm. b) Normalized fluorescent intensity of nanowells over time for control samples. Control 1: High concentration - O_2_ gas bubbled into the solution ([O_2_] = 15.6 mg/l), Control 2: Atmospheric conditions ([O2] = 6 mg/l), and Control 3: Low concentration-oxygen scavenger is added ([O_2_] = 0 mg/l). *Fluorescence intensities were normalized (to t=0) to allow direct comparison of fluorescent increases/decreases. (n=3).*

The results show that in the case of the sample with low initial oxygen concentration (0 mg/L), the intensity of fluorescence decreased over time. This is expected as Sodium sulfite scavenged the oxygen completely from the sample leading to a low concentration and a high initial fluorescent intensity of the sample. However, since PDMS is oxygen permeable, a small flux of oxygen from the PDMS into the well would increase the overall oxygen concentration, which quenches the flurophore leading to a reduction in its intensity. In the second case of the sample with high initial oxygen concentration (15.6 mg/L), the intensity of fluorescence increased over time. Oxygen bubbling produced high oxygen concentration, which quenched the fluorophore significantly, and the initial fluorescent intensity was very low. Over time some of the oxygen would diffuse out into the PDMS surrounding the sample leading to lower overall concentration in the well and higher intensity. The increase or decrease in fluorescent intensity over time under these extreme conditions was found to be significantly small demonstrating that PDMS is an effective material to separate the sample from the ambient environment for metabolic monitoring. Finally, in the case of the sample solution exposed to ambient atmospheric conditions, it was found that the fluorescent intensity was nearly the same over long durations of time, indicating that any change in intensity would in subsequent experiments only be due to consumption processes inherent to the presence of bacteria in the wells.

### 4.4. Growth of bacteria in nanowells

It is important to determine that the bacteria grow normally in the confines of the nanowells and their growth is not restricted due to depletion of nutrients. In order to test this, a long duration growth study was performed where GFP expressing *E.coli* (10^5^ cells/mL) in LB media with the fluorophore was dispensed into the nanowell array (whereby each well contained a 1nL volume) and observed over 12 hrs. The results as shown in figure 4 demonstrated significant growth of the bacteria with several rounds of division. At time t=0 (figure 4a) only a few isolated bacteria are observed. At time t = 12 hours (figure 4b) large colonies of bacteria originating from the initial single bacteria could be observed. It is interesting to note that the growth happens in a filamentous form as the *E.coli* divides along its length and since there is no disturbance in this particular microenvironment, the daughter cells were located right next to the mother cells. This experiment clearly shows that the volume of the nanowell is sufficiently large to allow growth and division of bacteria over a period of 12 hours at least.

**Figure 4.**
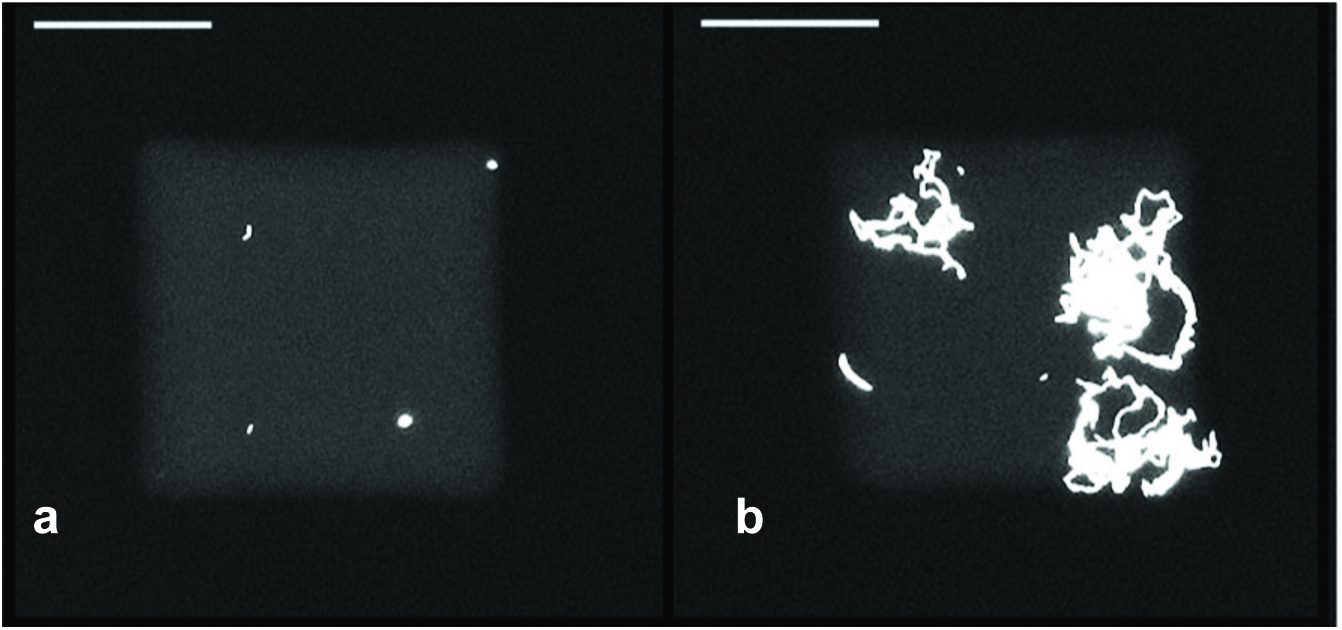
Fluorescent images of nanowells with GFP expressing bacteria *E.coli* K12 in LB media at different time points indicating growth of the bacteria in confined volumes. The well volume was 1 nL. a) Image at time t=0; b) image after 12 hrs of growth. Scale bars, 50 μm.

### 4.5. Measurement of metabolic activity

Fluorescent intensity corresponds to metabolic activity within the wells. Each bacterium consumes a certain amount of oxygen for its metabolism as long as it is viable. RTDP is oxygen quenching, so the fluorescent intensity increases with oxygen depletion within nanowells. Thus fluorescent intensity in the nanowells can be used as a marker for viability of bacteria that is present in that well. In order to test this principle, sample solutions containing 10^6^ cells/ml were loaded into a nanowell array with well volume of 1 nL and the fluorescent intensity in these wells imaged over time. Figure 5a shows the change in intensity between t=0 and t= 2.5 hrs. A clear increase in intensity was observed demonstrating that the metabolism of the bacteria in the solution can be observed using the intensity of the oxygen sensitive fluorophore in the nanowells. The intensity is also nearly uniform over the entire well indicating that a low resolution, low-cost imager can potentially be used for measuring the intensities. Next, in order to quantify the effect of bacterial concentration on the increase in intensity, sample solutions with various concentrations from 104 to 10^8^ cells/mL were dispensed into the wells and the change in intensity measure over time. Each concentration was repeated three times and in each case three wells were measured *(n=3).* The results are presented in figure 5b and show that in all cases, an increase in intensity occurs over time due to the consumption of oxygen in the wells by the bacteria present in them. Higher concentrations have a greater rate of increase in intensity indicating that the rate of consumption of oxygen was higher which is expected. These experiments indicate that the oxygen levels in the nanowells can be measured with an oxygen sensitive fluorophore RTDP that has high sensitivity and low cytotoxicity, to monitor the metabolic activity of bacteria as well as quantify their concentration in sample solution.

### 4.6. Drug effectiveness

In many cases of bacterial infection such as Tuberculosis, identification of drug effectiveness of the bacteria present in a particular sample quickly is of great importance to a physician in identifying the appropriate treatment.^39^ The effectiveness of any drug on that particular infection can be identified by adding it along with the growth medium and determining if the bacteria in the patient sample stops growing and loses its viability‥ Thus by adding the drug to the growth medium before dispensing it into the nanowells and measuring metabolic changes of the bacteria from its oxygen consumption, one could measure the effect of the drugs impact on the bacterial growth and survival. In order to test whether effect of a drug on bacterial metabolism could be identified in this device, an experiment was performed where ampicillin, which is a known drug that inhibits the growth of *E.coli* OP50 strain, was loaded along with it. In the experiment, 5 mg/L of ampicillin, a concentration that is well above the minimum inhibitory concentration (MIC) of 2mg/L^40^, was added into a bacterial broth with a concentration of 10^6^ cells per mL to inhibit the cell growth. The control was the same experiment without the drug loaded, and was repeated thrice *(n=3).* The profile of the increase in intensity with time is shown in figure 5c. It appears that in the devices where ampicillin was added, the fluorescent intensity did not change significantly, indicating that the oxygen in those nanowells was not consumed. Antibiotics like ampicillin act as an irreversible inhibitor of the enzyme transpeptidase, which is essential in OP50 *E.coli* bacteria to make their cell walls and thereby inhibits growth. If the bacteria were sensitive to the drug, there should be no (or little) evident increase in fluorescent intensity, which signifies that there is no metabolic activity. Damaged and dead bacterial cells have lower innate metabolic activity^41^, and thus will not consume oxygen. These results indicate that the ampicillin had the intended effect of preventing the growth and viability of the bacteria in the wells. The control on the other hand had a gradual and significant increase in intensity indicating that under the same growth condition and in the absence of ampicillin the bacteria were viable. These experiments indicate that metabolic monitoring of the nanowells can provide crucial information about the effectiveness of a particular drug on a particular infection rapidly. This will be crucial information for a physician deciding between various drugs available to treat patients with infections where drug resitance may be prevalent‥

**Figure 5.**
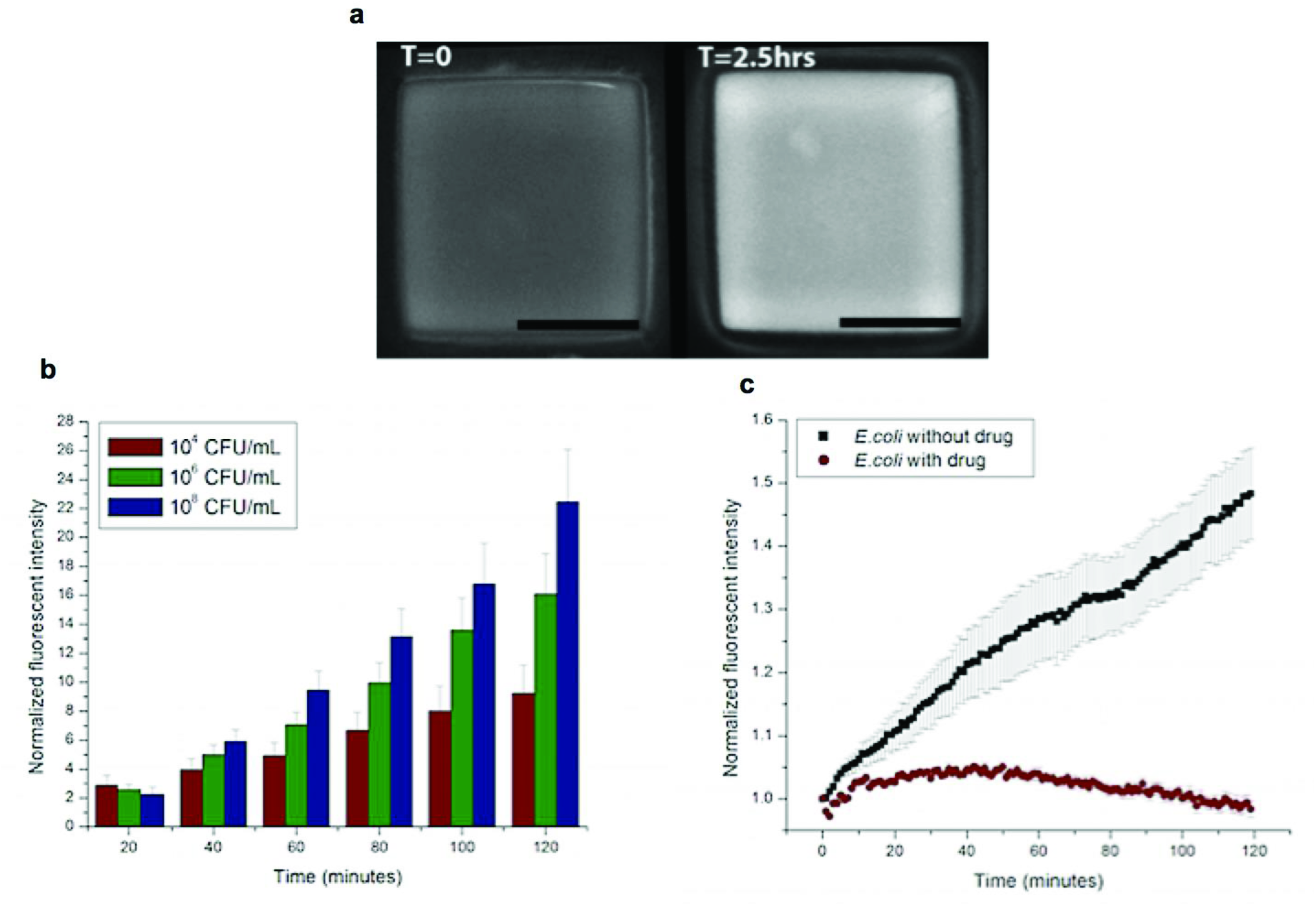
**a)**Fluorescence images of nanowell containing a sample solution with a bacterial concentration of 10^6^ CFU/mL at time t=0 and after 2.5 hrs showing a clear increase in fluorescent intensity over time which is indicative of oxygen depletion due to bacterial metabolism. Scale bars, 50 μm. **b)** Normalized fluoroscent intensity of the nanowells containing bacteria measured over time for various concentrations of bacteria. *Fluorescence intensities were normalized (to t=0) to allow direct comparison of fluorescent increase (n=3).* The devices used had a well size of 1 nL. Higher increases were observed when the concentration of bacteria in the wells were higher. **c)** Normalized fluoroscent intensity in nanowells measured over time in the presence and absence of ampicillin. The bacterial concentration used was 10^6^ CFU/ml. *Fluorescence intensities were normalized (to t=0) to allow direct comparison of fluorescent increases/decreases. (n=3)*

### 4.7. Effect of nanowell size

The size of the wells determine the amount of oxygen that is originally present in the wells and consequently, the time at which a bacteria in a well can consume sufficient amount of oxygen to reduce the concentration sufficently so that it is measurable. This effect of the nanowell size is likely to be noticeable at low concentrations of bacteria. In order to quantify this effect, sample solutions with *E.coli* (K12 strain) at a concentration of 10^4^ cells/mL were dispensed into devices with various well sizes of: 0.1 mL, 100 nL and 1nL respectively. Three devices at each well size were tested and three wells were measured in each device. The fluorescent intensity in these wells was measured over time and the results are plotted in figure 6. It shows that the device with smallest sized wells (1nL volume), had the largest increase in fluorescent intensity. In comparison, the device with 100nL wells showed a slower rate of increase and those with 0.1mL wells showed only a miniscule increase over time. At the concentration of 10^4^ cells/mL, the population density of the bacteria in the wells for the various well volumes are: 100's cells/well for 0.1 mL wells, ~1 cell/well for 100 nL and << 1 cell/well (or 1 well in a 100 are filled with one bacteria) for 1 nL wells. In the device with the 1 nL well, the effective concentration in those wells that have a bacteria is 1 cell/nL or 10^9^ cells/mL which is a 5 order of magnitude amplification in concentration due to segmentation of the sample. Therefore the bacteria in the nanowell can quickly deplete the oxygen present in the well and lead to faster detection. It also indicates that single cell detection is easily possible in this format.

**Figure 6.**
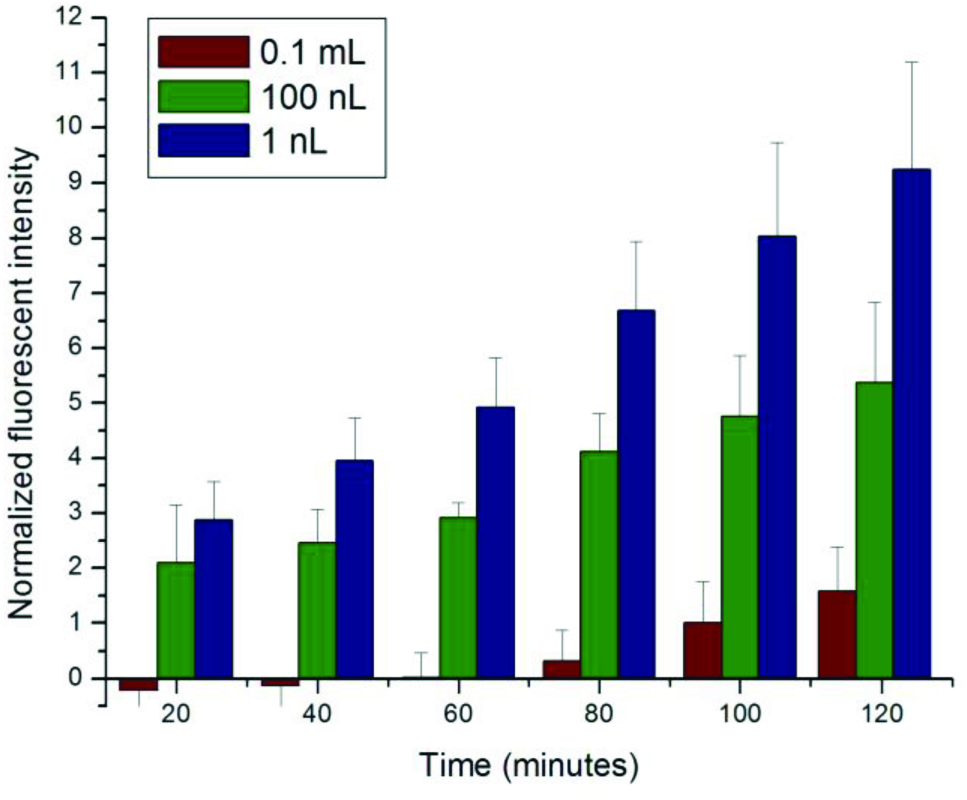
Fluoroscence intensity in nanowells measured over time for various well volumes. Fluorescence intensities were normalized (*to t=0*) to allow direct comparison of fluorescent increases (*n=3*). Smaller wells show a more significant intensity change corresponding to the rate of change of oxygen concentration in them.

## 5. CONCLUSION

Culture based methods are the gold standard for diagnosis of bacterial infection and provide crucial information about viability and drug effectiveness. As compared to molecular diagnostic methods, they are low cost, simple to use, do not requirement extensive sample preparation and therefore are more suitable for resource poor settings. However, they typically take 1 day to 4 weeks to obtain the results due to the dependence on growth and multiplication of the bacteria into colonies for identification. Instead, metabolic monitoring can be used to perform diagnosis of culture-based assays faster. In this paper we have developed a simple method to automatically aliquot a sample of various viscosities into thousands of nanoliter volumes and used it to measure metabolic activity of bacteria that are present in them. We demonstrate that the nanoscale volumes have sufficient nutrients to sustain the bacteria for more than 10 hours. We show that the metabolic rate can be easily measured in these wells by using an oxygen sensitive fluorophore and that the rate of change of its intensity is proportional to the concentration of the bacteria in the sample. We also show single bacterial cell detection is possible and that the time for detection is greatly reduced if the sample is aliquoated into thousands of small nanoliter wells and measured in parallel. Finally, we demonstrate that the same assay could be used to identify the effectiveness of a drug on that particular infection which provides crucial information to a medical professional in formulating a treatment. Apart from medical diagnosis, these nanowell devices for metabolic monitoring have many potential applications for many industries (i.e. medical and food) and to be able to test and study bacteria with rapid, accurate, and effective responses.

## 6. Acknowledgements

We acknowledge financial support from Grand Challenges Canada through their Rising Stars in Global Health Award, The Canada Research Chairs Program and the Ontario Research Fund, Research Excellence Program. We also acknowledge personnel in the Biophotonics facility, Micro Nanosystems center and Biointerfaces institute at McMaster for space and equipment for conducting this research.

